# More than Two Populations of Microtubules Comprise the Dynamic Mitotic Spindle

**DOI:** 10.1101/769752

**Authors:** Aaron R. Tipton, Gary J. Gorbsky

## Abstract

The microtubules of the mitotic spindle mediate chromosome alignment to the metaphase plate, then sister chromatid segregation to the spindle poles in anaphase. Previous analyses of spindle microtubule kinetics utilizing fluorescence dissipation after photoactivation described two main populations, a slow and a fast turnover population, and these were ascribed to reflect kinetochore versus non-kinetochore microtubules, respectively. Here, we test this categorization by disrupting kinetochores through depletion of the Ndc80 complex. In the absence of functional kinetochores, microtubule dynamics still exhibit slow and fast turnover populations, though the proportion of each population and the timings of turnover are altered. Importantly, the data obtained following Hec1/Ndc80 depletion suggests other sub-populations, in addition to kinetochore microtubules, contribute to the slow turnover population. Further manipulation of spindle microtubules revealed a complex landscape. Dissection of the dynamics of microtubule populations provides a greater understanding of mitotic spindle kinetics and insight into roles in facilitating chromosome attachment, movement, and segregation during mitosis.

## Introduction

The mitotic spindle is a highly dynamic, spatially organized array of microtubules and associated proteins that orchestrates equal distribution of the chromosomes to daughter cells. Microtubules comprising the mammalian mitotic spindle at metaphase are often broadly classified into three categories. Kinetochore fiber microtubules, heterogeneous in length, together comprise a dense bundle that extend from poles to kinetochores with many making direct contact with kinetochores at their plus ends. Kinetochore fiber microtubules play major roles in chromosome movement and in regulating the spindle checkpoint through interactions with kinetochores. Astral microtubules emanate from spindle poles with many extending toward the cell cortex where they are important for spindle positioning. Interpolar microtubules extend from the poles to interdigitate at the spindle equator and play critical roles in spindle structure. Early examination of internal spindle microtubule dynamics through fluorescence dissipation after photoactivation assays demonstrated that dissipation data were best fit by a double exponential curve (Zhai et al., 1995). Two populations of microtubules were detected, a fast turnover and a slow turnover population. These two populations were equated with non-kinetochore and kinetochore microtubules respectively, which have been widely reported in early electron microscopic studies (Mastronarde et al., 1993; McIntosh et al., 1975; Rieder, 1981a; Rieder, 1981b; Rieder, 1982; Salmon et al., 1976; Wise et al., 1991). However, judging from ultrastructure, spindle microtubules likely include more than two populations. For example, electron microscopy identifies within kinetochore fibers microtubules that reach from pole to kinetochore, others linked to either the pole or kinetochore with the other end free, and microtubules with two free ends (McDonald et al., 1992; Rieder, 1981b). Potentially, all may exhibit different dynamics within the fiber. Additionally, microtubules that branch from kinetochore fibers and microtubules bridging sister kinetochores have been observed within the mitotic spindle (Kajtez et al., 2016; Kamasaki et al., 2013; Mastronarde et al., 1993; Petry et al., 2013). To better evaluate the multiplicity of microtubule turnover within metaphase mitotic spindles, we carried out photoactivation experiments while manipulating populations of microtubules.

## Results and Discussion

### Mitotic spindles in cells lacking functional kinetochores still reveal fast and slow turnover microtubule populations

Early fluorescence dissipation after photoactivation observations of spindle microtubule dynamics suggested the slow turnover population of microtubules within the spindle was comprised of kinetochore fiber microtubules based on observations demonstrating they are more stable than non-kinetochore microtubules (Cassimeris et al., 1990; Gorbsky and Borisy, 1989; Mitchison, 1988; Rieder, 1981b; Salmon et al., 1976; Salmon et al., 1984; Zhai et al., 1995). The Ndc80 complex, comprised of Hec1/Ndc80, Nuf2, Spc24, and Spc25 is the primary end-on microtubule binding component at kinetochores (DeLuca et al., 2005; DeLuca et al., 2006; McCleland et al., 2004; Wei et al., 2007). Therefore, if only two populations, kinetochore and non-kinetochore microtubules, exist within the central spindle of metaphase cells, disruption of the Ndc80 complex should eliminate the slow turnover population.

To test this, we measured fluorescence dissipation after photoactivation of microtubules in U2OS cells stably expressing photoactivatable GFP (PAGFP)-tubulin and mCherry-Tubulin. Cells transfected with either control or Hec1 siRNA, to prevent kinetochore-microtubule interaction, were analyzed. The proteasome inhibitor MG132 at 10 μM was included to prevent mitotic exit, since the spindle checkpoint is abrogated by strong inhibition of the Ndc80 complex (McCleland et al., 2003). Only cells entering mitosis during the time frame of imaging were used because we determined that turnover of microtubules is significantly slowed by prolonged metaphase arrest that occurs in cells treated with MG132 (Fig. S1), but not by brief arrest (see below). Mitotic spindles were located by mCherry-Tubulin fluorescence. A bar-shaped region within the spindle on one side of the metaphase plate was photoactivated, and time-lapse images were acquired. Representative images depict fluorescence in pre- and post-photoactivated cells and the mid-volume mCherry-Tubulin plane (Fig. 1, Top). As determined in previous photoactivation experiments with spindle microtubules (Zhai et al., 1995) data from control cells were best fit by double exponential curves with the formula

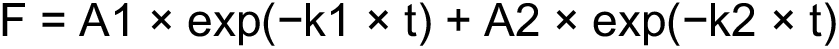

where A1 and A2 represent the percent total fluorescence contribution of the fast turnover and slow turnover microtubule populations, and k1 and k2 represent the respective decay rate constants (Fig. 1, bottom). However, even though Hec1 depletion largely eliminates end-on microtubule attachment at kinetochores, photoactivation data remained best fit by the double exponential suggesting that photoactivation still revealed two major populations of microtubules (Fig. 1, bottom). Fitting to single and triple exponential curves resulted in poorer R^2^ values (R^2^ < 0.990). In all instances, the data were best fit by a double exponential curve whose R^2^ value is reported in the figures. Successful Hec1 depletion was confirmed by comparing fluorescent DNA morphology to known Hec1 depletion phenotypes and by western blot analysis (Fig. S2) (DeLuca et al., 2005; Martin-Lluesma et al., 2002).

**Figure 1.**
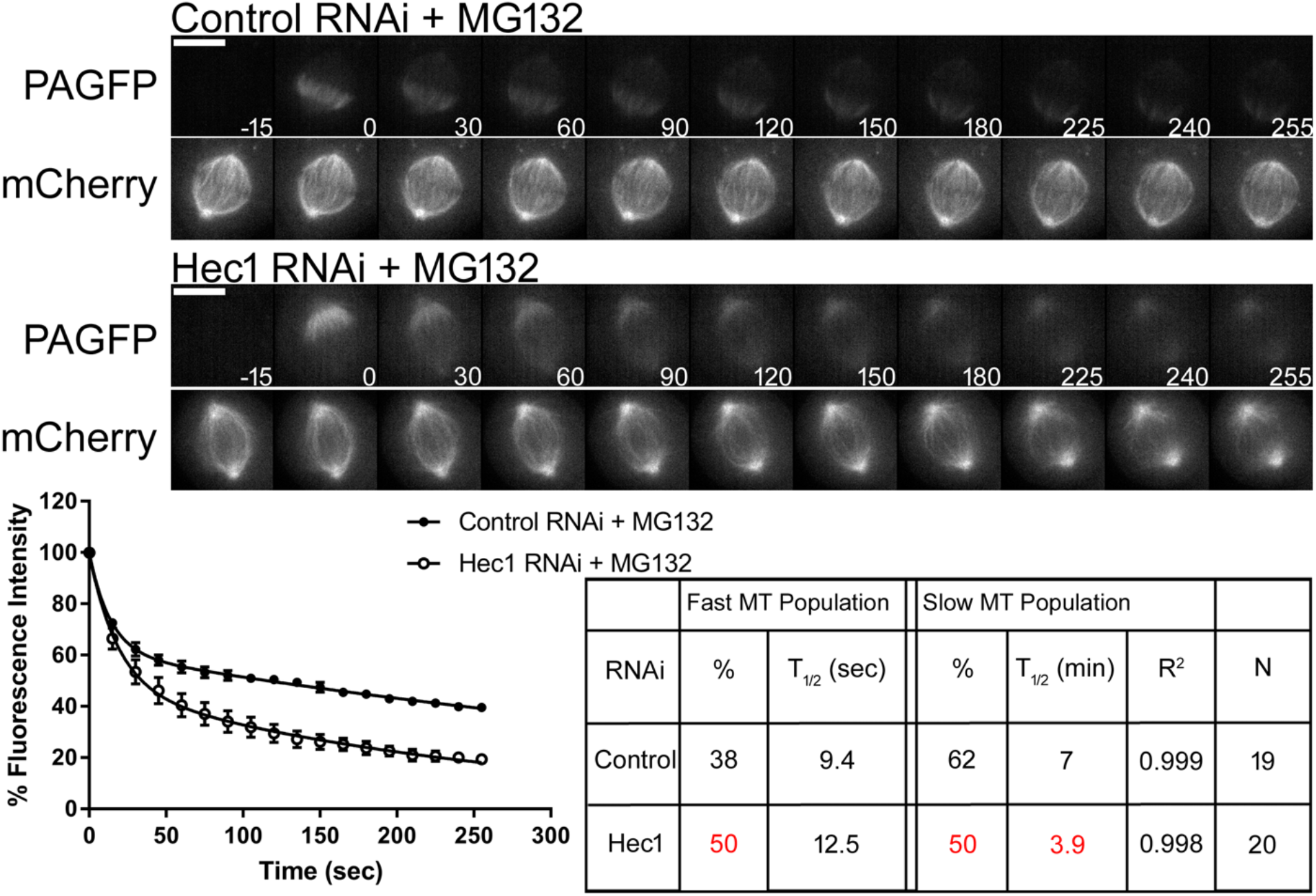
End-on attached microtubules comprise only a portion of the slow turnover microtubule population. (Top) Select frames from live cell imaging of Tubulin photoactivation in metaphase U2OS cells stably expressing photoactivatable GFP-Tubulin and mCherry-Tubulin following transfection with either control or Hec1 siRNA along with 10 μm MG132 treatment to prevent mitotic exit. mCherry-Tubulin frames are from the mid-volume plane. Only cells entering mitosis during imaging were photoactivated to ensure Tubulin turnover was not affected by prolonged mitotic arrest. Control cells were imaged at metaphase. Time, sec. Bar, 10 μm. (Bottom) Fluorescence dissipation after photoactivation. The filled and unfilled circles represent the average values recorded at each time point after photoactivation. The bars represent SEM. Control n = 19 cells, Hec1 n = 20 cells from three independent experiments. Lines indicate fitted curves (Control RNAi R^2^ = 0.999; Hec1 RNAi R^2^ = 0.998). Significant differences from control indicated in red in table.

While the total fluorescence contribution of the fast and slow turnover microtubule populations in control cells was measured to be 38% ± 2.4% and 62% ± 2.3%, respectively, cells depleted of Hec1 had both a significant increase in the fast turnover population to 50% ± 4.2% and decrease in the slow turnover population to 50% ± 4.2%. Thus, in Hec1-depleted cells a portion of the slow turnover microtubule population is reduced, and the fast turnover population increased. This suggests that mitotic spindles in cells unable to form end-on microtubule attachments are still comprised of multiple populations, fast turnover and slow turnover. When microtubule half-lives were examined, minor changes were observed between control and Hec1-depleted cells in the fast turnover population. In contrast, the *t*_1/2_ of the slow turnover population was significantly faster after Hec1 depletion (*t*_1/2_ = 3.9 ± 0.8 m) when compared with controls (*t*_1/2_ = 7 ± 0.6 m) (Fig. 1, bottom). These data reveal the existence of another slow population that does not include kinetochore fiber microtubules with end-on attachments to kinetochores. Thus contrary to the current assumptions regarding interpretation of photoactivation data the mitotic spindle includes one or more populations with relatively slow turnover that are not end-on kinetochore bound. These might include microtubules stabilized by lateral interactions with kinetochores, by association with chromosome arms, or by parallel or antiparallel bundling.

### Slow microtubule population turnover accelerates following loss of spindle bipolarity

Interpolar microtubules of opposite polarity that interdigitate plus ends near the spindle midline are another category of microtubules that comprise the mitotic spindle. Early classification of mitotic spindle structure suggested that most non-kinetochore microtubules were interpolar microtubules, concluding that interpolar microtubules constituted the bulk of the mitotic spindle (Mastronarde et al., 1993; McIntosh et al., 1975; Rieder, 1981a). Later, the fast turnover population of microtubules within the mitotic spindle was inferred to be comprised of interpolar microtubules (Zhai et al., 1995). To eliminate interdigitating, interpolar microtubules, cells were treated with the Eg5 inhibitor S-trityl-L-cysteine (STLC) (10 μM) to block bipolar spindle formation, leaving microtubules extending from the unseparated centrosomes with uniform polarity, some of which are free in the cytoplasm while others interact with kinetochores or with chromosome arms. Cells were analyzed as described in Fig. 1. Representative images depict fluorescence in pre- and post-photoactivated cells and the mid-volume fluorescence mCherry-Tubulin plane (Fig. 2, Top). While the total fluorescence contribution of the fast and slow turnover microtubule populations in control cells was measured to be 33% ± 3.4% and 67% ± 3.5%, respectively, cells treated with STLC had both a minor increase in the fast turnover microtubule population to 42% ± 3.5% and a minor decrease in the slow turnover microtubule population to 58% ± 3.4%. Thus, in cells treated with STLC, a portion of the slow turnover microtubule population is reduced, resulting in an increase in fast turnover population. In cells prevented from forming bipolar spindles (lacking interdigitating interpolar microtubules), microtubules still include fast and slow turnover populations. When microtubule half-lives were examined, no significant changes were observed between control and STLC treated cells in the fast turnover population. In contrast, the *t*_1/2_ of the slow turnover population was significantly faster following STLC treatment (*t*_1/2_ = 3.3 ± 0.3 m) when compared with controls (*t*_1/2_ = 5.7 ± 0.5 m) (Fig. 2, Bottom). Contrary to previous assumptions, the data are consistent with the interpretation that interdigitating interpolar microtubules may also constitute a portion of the slow turnover population within the mitotic spindle. However, we cannot rule out that the faster *t*_1/2_ of the slow turnover population may reflect changes in dynamic microtubules attached to kinetochores lacking bipolar attachment and tension, resulting in decreased stability. Similarly, increases in the proportion of astral microtubules may account for the minor increase in the fast turnover population.

**Figure 2.**
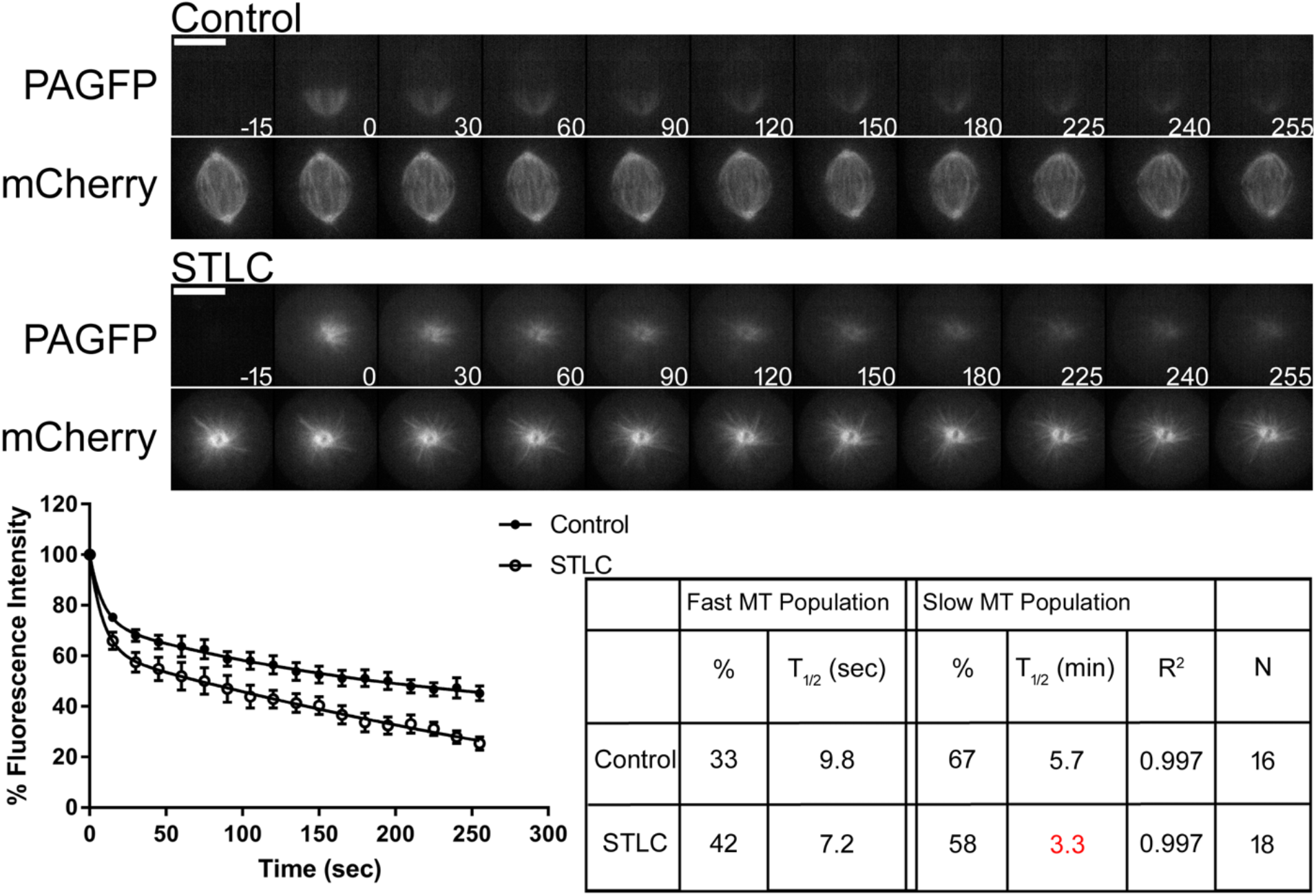
Interpolar microtubules also comprise a portion of the slow turnover microtubule population. (Top) Select frames from live cell imaging of Tubulin photoactivation in metaphase U2OS cells stably expressing photoactivatable GFP-Tubulin and mCherry-Tubulin following treatment with either control DMSO or 10 μm STLC to induce monopolar spindles. Cells were incubated 30 min prior to imaging. mCherry-Tubulin frames are from the mid-volume plane. Only cells entering mitosis during imaging were photoactivated to ensure Tubulin turnover was not affected by prolonged mitotic arrest. Control cells were imaged at metaphase. Time, sec. Bar, 10 μm. (Bottom) Fluorescence dissipation after photoactivation. The filled and unfilled circles represent the average values recorded at each time point after photoactivation. The bars represent SEM. Control n = 16 cells, STLC n = 18 cells from four independent experiments. Lines indicate fitted curves (Control R^2^ = 0.997; STLC R^2^ = 0.997). Significant differences from control indicated in red in table.

We next sought to displace Eg5 from metaphase arrested cells to determine if a subset of interpolar microtubules are lost or if other factors function to stabilize these microtubules resulting in altered dynamic properties. To accomplish this cells were arrested at metaphase with MG132 then subsequently treated with DMSO or STLC. Consistent with previous observations in other cell lines, U2OS cells maintain biopolar spindles under these conditions indicating that many cell types do not require Eg5 for maintenance of metaphase spindle bipolarity (Gayek and Ohi, 2014; Kollu et al., 2009). We did not observe a statistical difference in fast or slow microtubule *t*_1/2_ between control MG132 arrested cells and MG132 arrested cells treated with STLC (Fig. S3).

Unfortunately our efforts to simultaneously reduce both kinetochore-attached microtubules and interdigitating, interpolar microtubules were unsuccessful. Hec1-depleted cells treated with 100 μM Monastrol formed bipolar spindles. Representative images depict fluorescence in pre- and post-photoactivated cells and the mid-volume fluorescence mCherry-Tubulin and DNA plane (Fig. S4 A). This finding suggests that functional end-on kinetochore attachments are required to collapse spindles in mitotic cells treated with Eg5 inhibitors. However, in a previous study, siRNA was used to deplete Nuf2, another component of the Ndc80 complex, and there, co-treatment with Eg5 inhibitor induced spindle collapse and formation of monopolar spindles (Ganem and Compton, 2004). We hypothesized that in the previous study, incomplete Nuf2 depletion may have left enough intact Ndc80 complex to produce partially functional end-on attached kinetochores. To test this hypothesis, we treated cells with low (20 nM), or high (100 nM) concentrations of Nuf2 siRNA. These cells were treated with 100 μM Monastrol for 1 h, fixed and labeled for Nuf2, α-Tubulin, and ACA by immunofluorescence. Cells were then scored for monopolar and bipolar spindles in each condition (Fig. S4 B, left). In cells treated with the low concentration of Nuf2 siRNA, the majority (65%) were found to contain monopolar spindles. In contrast, in cells treated with the high concentration of Nuf2 siRNA, a minority (23%) exhibited monopolar spindles. The minority population forming monopolar spindles in cells treated with the high concentration of Nuf2 siRNA displayed detectable levels of Nuf2 at kinetochores indicating incomplete knockdown (Fig. S4 B). Live cell imaging of cells entering mitosis following low and high Nuf2 siRNA confirmed the above results (Movie S1, S2). These data are consistent with our initial observation (Fig. S4 A) and suggests that our Hec1 depletion more effectively eliminates microtubule attachment to kinetochores than does Nuf2 depletion. Our results are also consistent with the biphasic effects of Ndc80 complex inactivation on the spindle checkpoint. Partial inactivation of the complex induces a spindle checkpoint-dependent arrest, suggesting residual intact Ndc80 complex at kinetochores, while complete inactivation of Ndc80 complex function eliminates spindle checkpoint signaling causing cells to prematurely exit mitosis (Martin-Lluesma et al., 2002; McCleland et al., 2003).

We attempted to quantify spindle microtubule dynamics in monopolar spindles by inhibiting Eg5 in cells with partially reduced Ndc80 activity after treatment with the low concentration of Nuf2 siRNA. However, the fluorescence dissipation after photoactivation data obtained from these experiments resulted in extremely poor regression curve fits rendering the data unusable (data not shown). We surmised that the partial inactivation of the Ndc80 complex led to variable effects on intact end-on kinetochores causing erratic variations in microtubule dynamics.

### Turnover of both fast and slow microtubule populations is slower following temperature reduction

A well-established characteristic of spindle microtubules in cell culture models is the sensitivity of stability to changes in temperature (Brinkley and Cartwright, 1975; Rieder, 1981a; Rieder, 1981b; Zhai et al., 1995). In the LLC-PK cell line, decreases in temperature resulted in increases in the *t*_1/2_ of the slow turnover microtubule population, with no major effect on the *t*_1/2_ of the fast turnover population (Zhai et al., 1995). This study also reported an overall decrease in the fluorescence contribution of the fast turnover population and increase of the slow turnover population following reductions in temperature. We sought to examine the effects of reductions in temperature on spindle microtubule dynamics in the U2OS cell line. Cells were imaged at 37°C (Control) as described above or at room temperature (22°C-25°C) (Low Temperature) as described in methods section. Representative images depict fluorescence in pre- and post-photoactivated cells and the mid-volume fluorescence mCherry-Tubulin plane (Fig. 3, Top). In contrast to the report from Zhai et al. (Zhai et al., 1995), we detected non-significant differences in the fluorescence contribution of fast and slow turnover microtubule populations when comparing control cells to cells imaged at low temperature. Major increases in *t*_1/2_, indicating slower turnover, were observed in cells imaged at low temperature in both the fast turnover population (Control *t*_1/2_ = 7.1 ± 0.6 s; Low Temperature *t*_1/2_ = 15.7 ± 1.2 s) and the slow turnover population (Control *t*_1/2_ = 4.7 ± 0.5 m; Low Temperature *t*_1/2_ = 14 ± 1.1 m) (Fig. 3, Bottom). This suggests that the *t*_1/2_ of both populations detected are sensitive to reduction of temperature. This is in contrast to the observations made by Zhai et al. (Zhai et al., 1995), which only reported a slower *t*_1/2_ in the slow turnover microtubule population. One potential explanation for the discrepancies could be differences in cell type. Additionally, advances in imaging technology may have permitted us to achieve higher temporal resolution allowing detection of differences in *t*_1/2_ of the fast turnover microtubule population.

**Figure 3.**
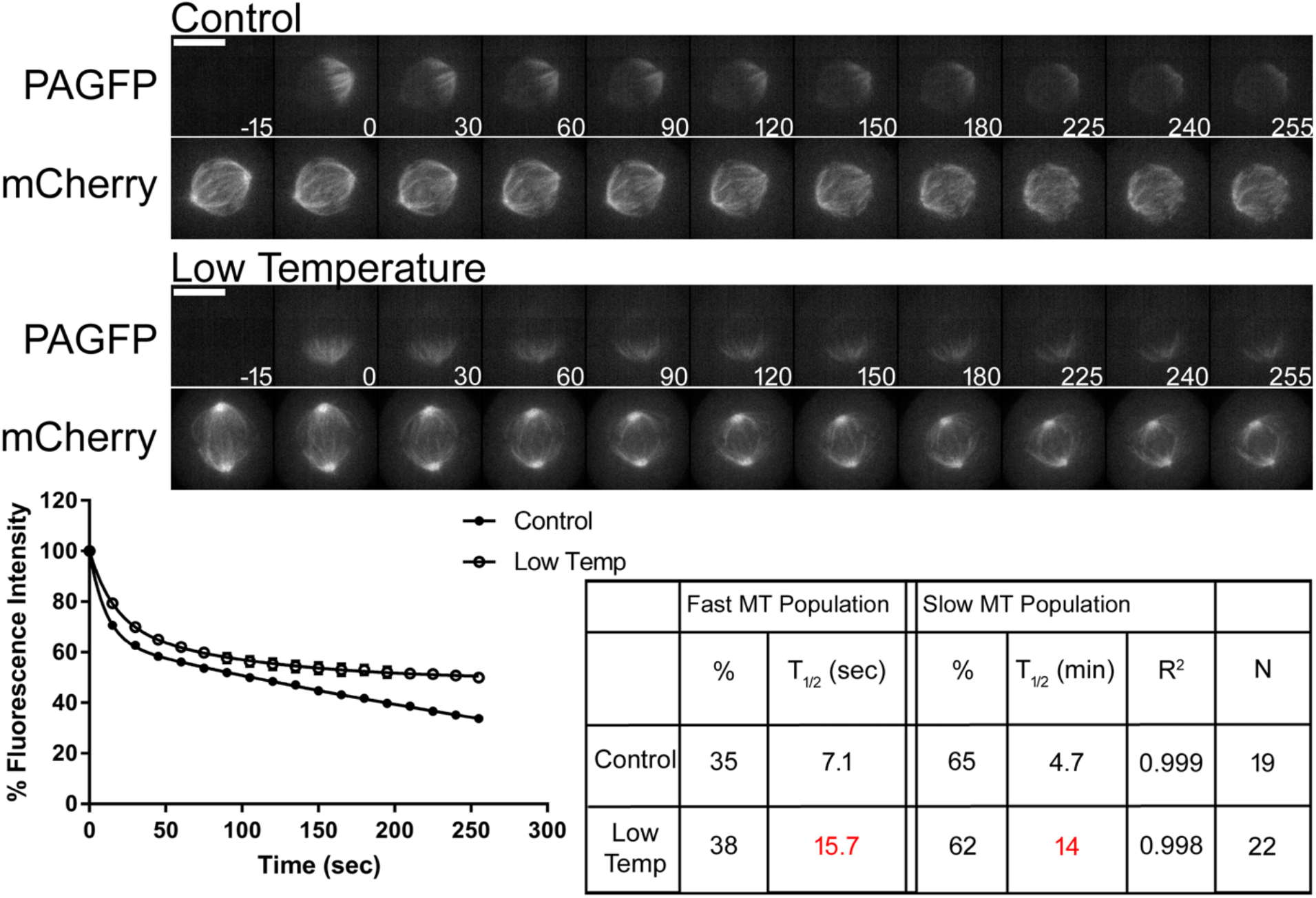
Both fast turnover and slow turnover microtubule populations are sensitive to decreases in temperature. (Top) Select frames from live cell imaging of Tubulin photoactivation in metaphase U2OS cells stably expressing photoactivatable GFP-Tubulin and mCherry-Tubulin following imaging at 37°C (standard conditions) or at low temperature (22°C-25°C) without objective heater, air curtain, and media warmed to room temperature. mCherry-Tubulin frames are from the mid-volume plane. Only cells entering mitosis during imaging were photoactivated to ensure Tubulin turnover was not affected by prolonged mitotic arrest. Cells were imaged at metaphase. Time, sec. Bar, 10 μm. (Bottom) Fluorescence dissipation after photoactivation. The filled and unfilled circles represent the average values recorded at each time point after photoactivation. The bars represent SEM. Control n = 19 cells, Low Temperature n = 22 cells from four independent experiments. Lines indicate fitted curves (Control R^2^ = 0.999; Low Temperature R^2^ = 0.998). Significant differences from control indicated in red in table.

### Turnover of the slow microtubule population is faster following Aurora B kinase inhibition

Aurora B kinase is an important regulator of kinetochore-microtubule attachment (Cheeseman et al., 2002; Cimini et al., 2006; Murata-Hori and Wang, 2002; Pinsky et al., 2006; Welburn et al., 2010). Aurora B responds to improper microtubule attachment to kinetochores by phosphorylating the N-terminal tail of Hec1/Ndc80, which reduces microtubule binding affinity. A previous report demonstrated in the PtK1 cell line that inhibition of Aurora B kinase activity via the small molecule ZM447439 caused an extensive increase (over seven-fold) in the *t*_1/2_ of the slow turnover microtubule population, with no major effect on the *t*_1/2_ of the fast turnover population when compared to control metaphase cells (Cimini et al., 2006). We in turn sought to examine the effects of Aurora B kinase inhibition on spindle microtubule dynamics in the U2OS cell line. To inhibit Aurora B kinase activity cells were treated with either 3 μM ZM447439 or DMSO (control) along with 10 μM MG132 to prevent mitotic exit. Cells were then analyzed as described above. Representative images depict fluorescence in pre- and post-photoactivated cells and the mid-volume fluorescence mCherry-Tubulin plane (Fig. 4, Top). No significant changes in the total fluorescence contribution of fast and slow turnover populations were observed between control cells and cells treated with ZM447439. When microtubule *t*_1/2_ was examined, no significant changes were observed between control and ZM447439-treated cells in the fast turnover population. In contrast, the *t*_1/2_ of the slow turnover population was significantly slower following ZM447439 treatment (*t*_1/2_ = 13.9 ± 0.7 m) when compared with controls (*t*_1/2_ = 8.5 ± 1.5 m) (Fig. 4, Bottom).

**Figure 4.**
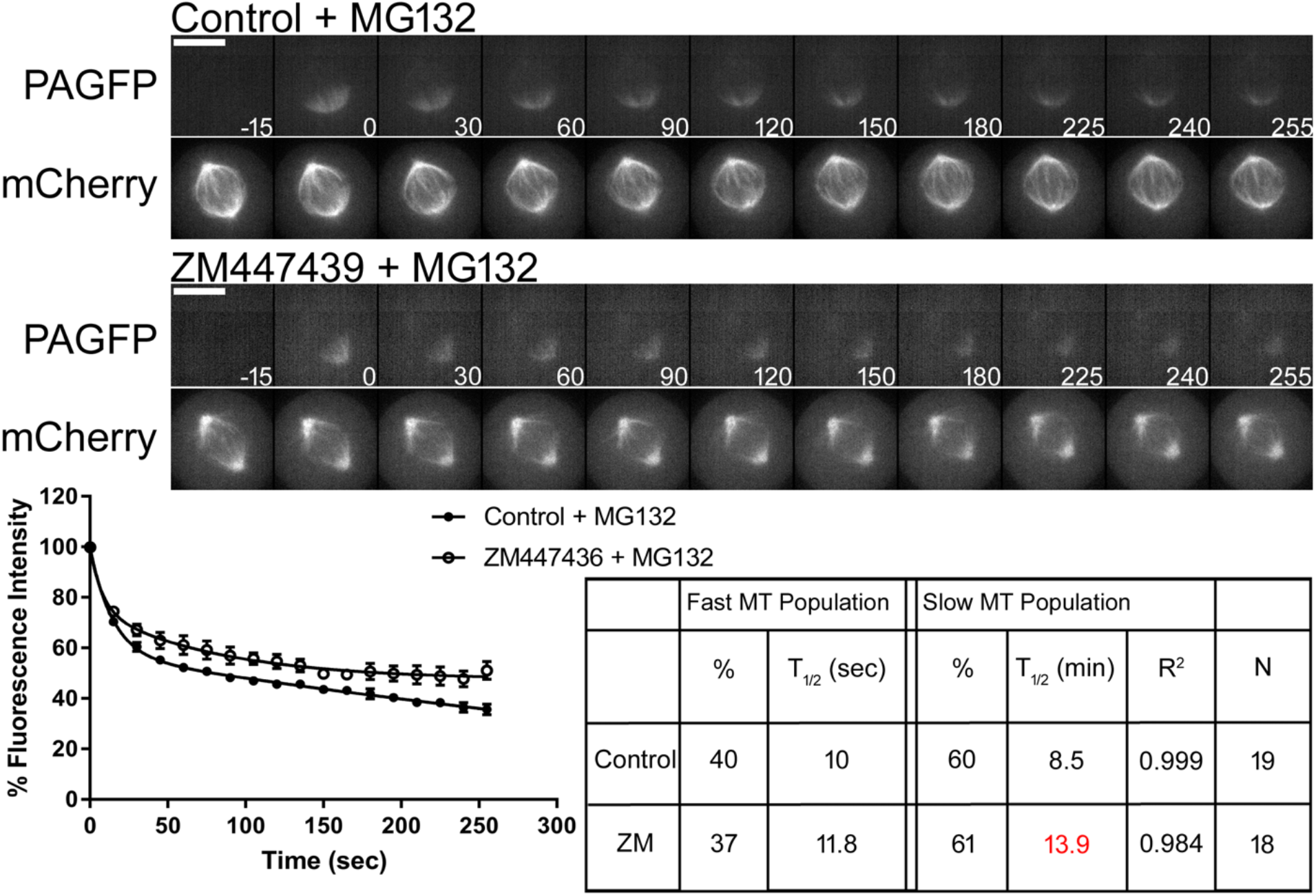
Aurora B kinase inhibition affects slow microtubule population turnover. (Top) Select frames from live cell imaging of Tubulin photoactivation in metaphase U2OS cells stably expressing photoactivatable GFP-Tubulin and mCherry-Tubulin following treatment with either control DMSO or 3 μm ZM447439 to inhibit Aurora B activity along with 10 μm MG132 treatment to prevent mitotic exit. Cells were incubated 30 min prior to imaging. mCherry-Tubulin frames are from the mid-volume plane. Only cells entering mitosis during imaging were photoactivated to ensure Tubulin turnover was not affected by prolonged mitotic arrest. Control cells were imaged at metaphase. Time, sec. Bar, 10 μm. (Bottom) Fluorescence dissipation after photoactivation. The filled and unfilled circles represent the average values recorded at each time point after photoactivation. The bars represent SEM. Control n = 19 cells, ZM447439 n = 18 cells from three independent experiments. Lines indicate fitted curves (Control R^2^ = 0.999; ZM447439 R^2^ = 0.984). Significant differences from control indicated in red in table.

### Aurora B kinase stabilizes non end-on attached slow turnover microtubules

To further examine the role Aurora B kinase activity plays in regulating spindle microtubule dynamics during mitosis, Aurora B kinase activity was inhibited in cells depleted of Hec1/Ndc80. We hypothesized that if Aurora B solely regulated end-on attached kinetochore microtubules, then Aurora B inhibition should not alter spindle microtubule dynamics in cells depleted of functional kinetochores. To test this hypothesis, cells were transfected with control siRNA followed by DMSO treatment or Hec1 siRNA followed by 3 μM ZM447439 treatment and analyzed as described above. 10 μM MG132 was included to prevent mitotic exit. Representative images depict fluorescence in pre- and post-photoactivated cells and the mid-volume fluorescence mCherry-Tubulin plane (Fig. 5, Top). We did not detect significant changes in the portion of the fast and slow turnover populations when comparing control-depleted and Hec1-depleted cells treated with Aurora kinase inhibitor. When microtubule *t*_1/2_ were examined, no significant changes were observed between control cells and cells depleted of Hec1 followed by ZM447439 treatment in the fast turnover population Strikingly however, the *t*_1/2_ of the slow turnover population was significantly faster in cells depleted of Hec1 followed by ZM447439 treatment (*t*_1/2_ = 2.2 ± 0.3 m) when compared with controls (*t*_1/2_ = 7.8 ± 1.3 m) (Fig. 5, Bottom). The *t*_1/2_ of the slow turnover population in cells depleted of Hec1 followed by ZM447439 treatment was also faster when compared to cells depleted of Hec1 alone (Hec1 RNAi *t*_1/2_ = 3.3 ± 0.3 m; Hec1 RNAi + ZM447439 *t*_1/2_ = 2.2 ± 0.3 m). Together these data suggest that Aurora B kinase plays roles in both stabilizing and destabilizing slow turnover microtubules within the mitotic spindle (comparing ZM447439 treatment alone and Hec1 depletion followed by ZM447439 treatment to their respective controls) (Fig. S5). One potential mechanism by which Aurora B kinase plays a role in stabilizing slow turnover microtubules is via regulation of localization and activity of microtubule depolymerases, such as the kinesin 13 family (Andrews et al., 2004; Bakhoum et al., 2009; Knowlton et al., 2006; Knowlton et al., 2009; Lan et al., 2004; Ohi et al., 2003; Ohi et al., 2004; Sampath et al., 2004). However, we cannot rule out other potential mechanisms by which Aurora B kinase functions in stabilizing slow turnover microtubule populations. Figure S6 is a compilation of all data presented above with statistical analyses.

**Figure 5.**
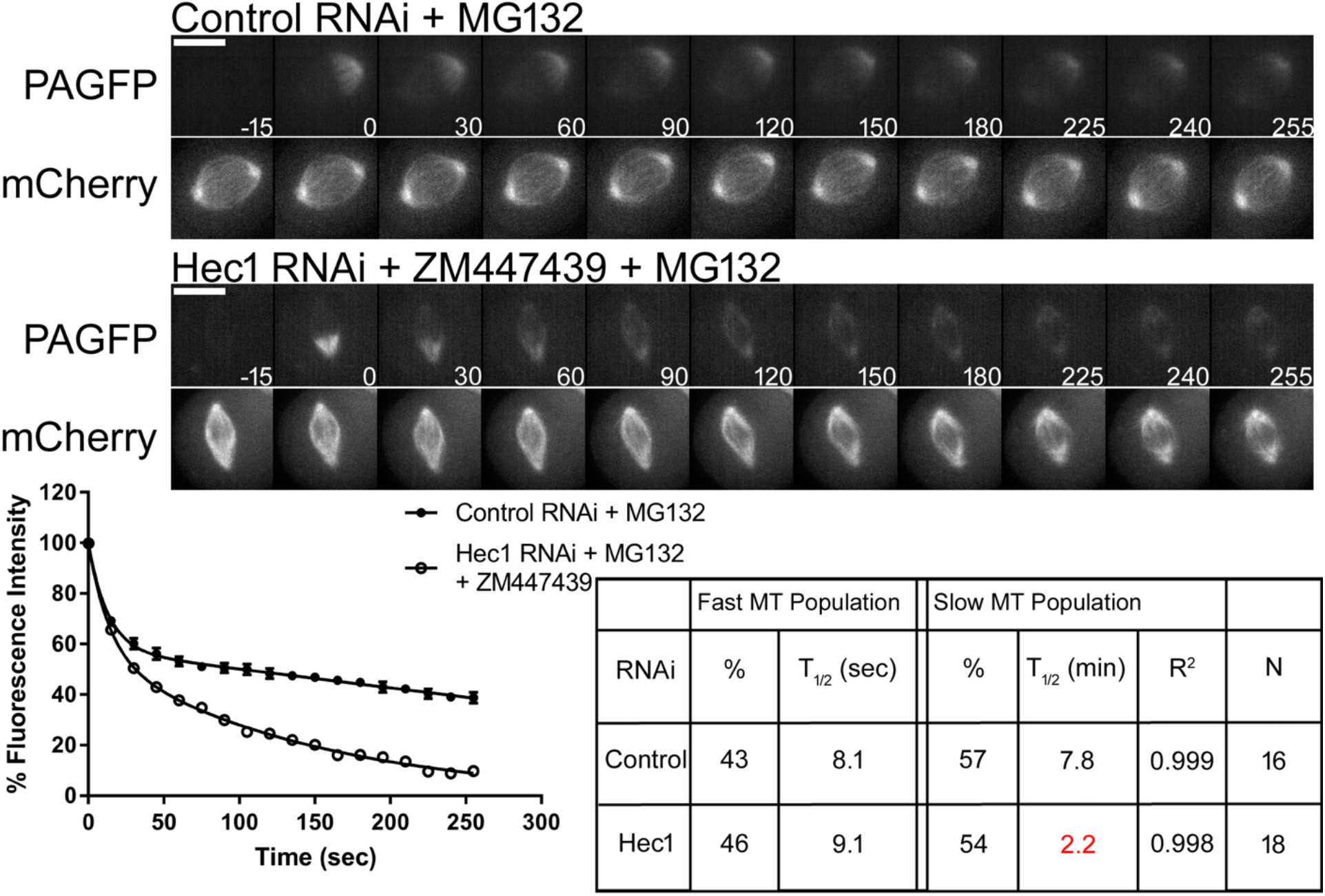
Aurora B kinase activity enhances stability of non-kinetochore slow turnover microtubules. (Top) Select frames from live cell imaging of Tubulin photoactivation in metaphase U2OS cells stably expressing photoactivatable GFP-Tubulin and mCherry-Tubulin following transfection with either control or Hec1 siRNA along with 10 μm MG132 treatment to prevent mitotic exit followed by control DMSO or 3 μm ZM447439 to inhibit Aurora B activity. Cells were incubated 30 min prior to imaging. mCherry-Tubulin frames are from the mid-volume plane. Only cells entering mitosis during imaging were photoactivated to ensure Tubulin turnover was not affected by prolonged mitotic arrest. Control cells were imaged at metaphase. Time, sec. Bar, 10 μm. (Bottom) Fluorescence dissipation after photoactivation. The filled and unfilled circles represent the average values recorded at each time point after photoactivation. The bars represent SEM. Control n = 16 cells, Hec1 + ZM447439 n = 18 cells from three independent experiments. Lines indicate fitted curves (Control RNAi R^2^ = 0.999; Hec1 RNAi + ZM447439 R^2^ = 0.998). Significant differences from control indicated in red in table.

The mitotic spindle is essential in the equal distribution of genetic material during mitosis. Early studies examining spindle microtubule dynamics determined the existence of two populations within the mitotic spindle (Zhai et al., 1995). A fast population that turns over in seconds and a slow population that turns over in minutes. These two populations have generally be equated with non-kinetochore microtubules (comprised of interdigitating interpolar microtubules) and kinetochore microtubules (comprised of microtubules attached to kinetochores), respectively. However, this nomenclature arose from observations indicating that microtubules attached to kinetochores displayed increased levels of stability while non-kinetochore microtubules represent the bulk of microtubules that constitute the mitotic spindle (Brinkley and Cartwright, 1975; Cassimeris et al., 1990; Gorbsky and Borisy, 1989; Mastronarde et al., 1993; McIntosh et al., 1975; Mitchison, 1988; Rieder, 1981a; Rieder, 1981b; Salmon et al., 1976; Salmon et al., 1984; Zhai et al., 1995). In this study, we sought to determine if this assumption reflected an oversimplification and if mitotic spindle dynamics could be tracked through various manipulations to reveal multiple subpopulations. We targeted end-on attached kinetochore fibers by depleting Hec1/Ndc80. In the absence of this major microtubule binding component at kinetochores, the slow turnover population was primarily affected resulting in both a decrease in fluorescence contribution and a faster *t*_1/2_ when compared to controls (Fig. 1). Data obtained following Hec1/Ndc80 depletion was best fit by a double exponential curve, indicating the existence of multiple populations in the absence of end-on attached kinetochore microtubules. This suggests that end-on attached kinetochore fibers are not the sole population comprising slow turnover microtubules.

To target interdigitating interpolar microtubules, Eg5 inhibition was used to prevent bipolar spindle formation and formation of interdigitating interpolar microtubules. In contrast to previous assumptions suggesting that interpolar microtubules comprise the non-kinetochore (or fast turnover) population, we found that STLC treatment both decreased the fluorescence contribution of the slow turnover population with a faster *t*_1/2_ when compared to controls (Fig. 2). However, we cannot rule out that the faster *t*_1/2_ of the slow turnover population or the decrease in the slow turnover population may in part be due to microtubules interacting with kinetochores lacking bipolar attachment/tension, or with microtubules associating with chromosome arms. Analysis of monopolar spindles likely includes free astral microtubules. Their addition to the rapid turnover population may result in a decrease in the slow turnover population. Additionally, we found Eg5 activity did not play a role in regulating spindle microtubule dynamics following the establishment of a bipolar spindle (Fig. S3).

When spindle microtubule dynamics were examined in conditions of reduced temperature, we found the *t*_1/2_ of both the fast and slow turnover populations to be slower, with little change in the relative fluorescence contribution (Fig. 3). This was in contrast to previous observations reporting changes in the *t*_1/2_ in the slow turnover population with no significant change in the *t*_1/2_ in the fast turnover population (Zhai et al., 1995). However, these discrepancies may have been a result of differences in cell types or advances in imaging technology.

The well-recognized regulator of kinetochore-microtubule attachment, Aurora B kinase, was also analyzed for its role in regulating microtubule dynamics within the mitotic spindle. In partial agreement with a previous report (Cimini et al., 2006), we found the *t*_1/2_ of the slow turnover population was slower following Aurora B kinase inhibition. However, the previous study, using the PtK1 cell line, reported extreme increases in stability for the slow turnover population with *t*_1/2_ of 52.6 min at 3 uM ZM447439 and 230.0 min at 20 uM, compared to 7.4 min for control metaphase cells (Cimini et al., 2006). Increased stability of the slow turnover population is consistent with the established role of Aurora B kinase in destabilizing microtubule-kinetochore attachments via phosphorylation of Hec1 (Cheeseman et al., 2002; Cimini et al., 2006; Murata-Hori and Wang, 2002; Pinsky et al., 2006; Welburn et al., 2010). In cells depleted of Hec1 and treated with Aurora B kinase inhibitor, the *t*_1/2_ of the slow turnover population was drastically faster. This suggests Aurora B kinase not only functions in destabilizing incorrect microtubule attachments to kinetochores, but also plays a role in enhancing microtubule stability of other populations within the mitotic spindle, perhaps through regulation microtubule depolymerases, such as those of the kinesin 13 family (Andrews et al., 2004; Bakhoum et al., 2009; Knowlton et al., 2006; Knowlton et al., 2009; Lan et al., 2004; Ohi et al., 2003; Ohi et al., 2004; Sampath et al., 2004).

Together, this study demonstrates that turnover of microtubules within the mitotic spindle is significantly more complex than previous models suggest. The mitotic spindle is likely comprised of multiple subpopulations. The identity of the fast turnover microtubule population within the mitotic spindle and mechanisms to manipulate its turnover (apart from reductions in temperature) remain elusive. Additionally, an inherent assumption made in this study is that spindle organization following various manipulations does not create new categories of microtubules. However, this may not be the case. For example in cells depleted of Hec1, non-K-fiber microtubules may assume an architecture that is not present in cells containing K-fibers. Testing this hypothesis will require more extensive manipulation of known microtubule associated proteins. Moreover, tubulin itself is also subject to post-translational modification (Barisic and Maiato, 2016; Barisic et al., 2015; Magiera and Janke, 2014; Song and Brady, 2015; Strzyz, 2016; Wloga et al., 2017; Yu et al., 2015). Examining how these various post-translational modifications play a role in regulating spindle microtubule dynamics during mitosis will be challenging but may lead important insights into mitotic spindle assembly and function.

## Materials and Methods

### siRNA

The siRNA sequence (5’-GAGUAGAACUAGAAUGUGA-3’) targeting Hec1 was synthesized by Qiagen. The ON-TARGETplus SMARTpool siRNA sequences (5’-GAACGAGUAACCACAAUUA-3’), (5’-UAGCUGAGAUUGUGAUUCA-3’), (5’-GGAUUGCAAUAAAGUUCAA-3’), and (5’-AAACGAUAGUGCUGCAAGA-3’) targeting Nuf2 was synthesized by Dharmacon. Non-targeting control siRNA was synthesized by Bioneer.

### Cell culture, transfection, and drug treatments

U2OS cells stably expressing photoactivatable GFP-Tubulin (PAGFP-Tubulin) and mCherry-Tubulin (see acknowledgments) were cultured in DMEM with 10% fetal bovine serum (FBS) supplemented with penicillin and streptomycin, 20mM HEPES, and 0.1 mM nonessential amino acids (NEAA) at 37°C with 5% CO_2_. For siRNA transfection, cells were transfected with 50 nM (Hec1) or 20 nM or 100 nM (Nuf2) siRNA for 48 h using Lipofectamine RNAiMAX (Invitrogen) according to the manufacturer’s protocol. To inhibit Aurora B activity, cells were treated with 3 μM ZM447439. 10 μM MG132 was also added to prevent mitotic exit. S-trityl-L-cysteine (STLC) or Monastrol was used at 10 μM and 100 μM respectively to induce monopolar spindles. Taxol was used at 10 μM. All drugs were added 30 min, except where indicated, prior to imaging. SiR-DNA (Cytoskeleton) was used at 500 nM to label DNA according to the manufacturer’s protocol.

### Photoactivation

PAGFP-Tubulin and mCherry-Tubulin expressing U2OS cells were grown on coverslips. To maintain appropriate pH levels and avoid evaporation during imaging, culture media was exchanged to Leibovitz’s L-15 medium supplemented with 10% FBS, penicillin, and streptomycin, and the medium was overlaid with mineral oil. Cells were treated as detailed in the figure legends and imaged using a 100x, NA 1.4 objective on a Zeiss Axio Observer inverted microscope equipped with an objective heater, air curtain, Yokogawa CSU-22 (Yokogawa) spinning disk, Mosaic (digital mirror device, Photonic Instruments/Andor), a Hamamatsu ORCA-Flash4.0LT (Hamamatsu Photonics), and Slidebook software (Intelligent Imaging Innovations). Photoactivation was achieved by targeting a selected area with filtered light from the HBO 100 via the Mosaic, and confocal GFP images (19 Z planes across 8 μm (0.5 μm/plane)) were acquired at 15 sec intervals for ~5 min. mCherry and SiR-DNA images were acquired at the mid volume Z plane. Only cells entering mitosis during imaging were photoactivated to ensure Tubulin turnover was not affected by prolonged mitotic arrest. To quantify fluorescence dissipation after photoactivation, we measured pixel intensities within an area surrounding the region of highest fluorescence intensity and background subtracted using an area from the nonactivated half spindle using MetaMorph software. The values were corrected for photobleaching by determining the percentage of fluorescence loss during image acquisition after photoactivation in the presence of 10 μM Taxol. Fluorescence values were normalized to the first time-point after photoactivation for each cell and the average intensity at each time point was fit to a double exponential curve F = A1 x exp(-k1 x t) + A2 x exp(-k2 x t), using SigmaPlot (SYSTAT Software), where A1 and A2 represent the fast turnover and slow turnover microtubule populations with decay rates of k1 and k2, respectively. t is the time after photoactivation. The turnover half-life for each population of microtubules was calculated as ln2/k. XY scatter plots were generated using GraphPad Prism. The Mann-Whitney two-tailed test in Prism (GraphPad Software) was used to determine statistical significance among groups.

### Immunofluorescence

PAGFP-Tubulin and mCherry-Tubulin expressing U2OS cells were grown on glass coverslips and treated as detailed in the figure legends. Cells were fixed in 2% paraformaldehyde/PHEM solution (60 mM PIPES, pH 6.9, 25 mM HEPES, 10 mM EGTA, 4 mM MgCl_2_) containing 0.5% Triton X-100 for 15 min. Coverslips were washed in MBST (10 mM MOPS, 150 mM NaCl, 0.05% Tween 20), blocked in 20% boiled normal goat serum (BNGS), and incubated overnight with primary antibodies. Samples were then incubated with secondary antibodies for 1 h, stained with DNA dye 4’,6-diamidino-2-phenylindole (DAPI) and mounted using Vectashield (Vector Laboratories). The following primary antibodies were used: Rat anti-α Tubulin (YL1-2), Rabbit anti-Nuf2 (P. Todd Stukenberg; University of Virginia) and human anti-Centromere Antibodies (ACA; Antibodies Incorporated). Secondary antibodies used were goat anti-rabbit antibodies conjugated fluorescein isothiocyanate (FITC; Jackson ImmunoResearch), goat anti-rat antibodies conjugated to Cy3 (Jackson ImmunoResearch), and goat anti-human antibodies conjugated to Cy5 (Jackson ImmunoResearch). The images were acquired using a Zeiss Axioplan II microscope equipped with a 100×, NA 1.4 objective and a Hamamatsu ORCA-ER camera (Hamamatsu Photonics) and processed using MetaMorph software. Quantification of immunofluorescence images was performed as previously described (Daum et al., 2009).

### Live-cell imaging

PAGFP-Tubulin and mCherry-Tubulin expressing U2OS cells were grown on glass coverslips. To maintain appropriate pH levels and avoid evaporation during imaging, culture media was exchanged to Leibovitz’s L-15 medium supplemented with 10% FBS, penicillin, and streptomycin, and the medium was overlaid with mineral oil. Cells were treated as detailed in figure legends. Time-lapse fluorescence images were collected using 100x, NA 1.4 objective on a Zeiss Axio Observer inverted microscope equipped with an objective heater, air curtain, Yokogawa CSU-22 (Yokogawa) spinning disk, Mosaic (digital mirror device, Photonic Instruments/Andor), a Hamamatsu ORCA-Flash4.0LT (Hamamatsu Photonics), and Slidebook software (Intelligent Imaging Innovations).. Images were captured every 3 min. Time-lapse videos displaying the elapsed time between consecutive frames were assembled using MetaMorph software.

### Western-blot analysis

For cell lysates, cells were arrested for 12 h with 100ng/ml Nocodozole and mitotic cells were collected then lysed in cell lysis buffer (1xPBS, 0.5% NP-40, 1 μM tris(2-carboxyethyl)phosphine, and 10% glycerol) supplemented with protease inhibitor cocktail (Sigma-Aldrich) and phosphatase inhibitors (100 mM NaF, 1 mM Na_3_VO_4_, 60 mM β-glycerophosphate, and 100 nM microcystin-LR). The protein concentration of lysates was measured using the BCA Protein Assay kit (Thermo Fisher Scientific). For electrophoresis, sample loading buffer (Invitrogen) and DTT to a final concentration of 50 mM were added. Proteins were separated with a NuPAGE gel electrophoresis system (Invitrogen) and transferred to a 0.45-μm PVDF membrane (Immobilon-FL; EMD Millipore). Membranes were blocked in 10% SEA BLOCK blocking buffer (Thermo Fisher Scientific) and 0.05% Tween-20 in 1xPBS (PBST). The following primary antibodies were used: rabbit anti-Hec1 (P. Todd Stukenberg; University of Virginia), mouse anti–actin (Abcam). Membranes were washed in PBST. Secondary goat anti–mouse and goat anti–rabbit antibodies were conjugated to either IR700 or IR800 dyes (Azure Biosystems). Western blots were imaged on an Azure c600 Imaging System.

## Abbreviations used

PAGFP: photoactivatable green fluorescent protein
RNAi: RNA interference
STLC: S-trityl-L-cysteine

## Acknowledgments

We thank, Marvin Tanenbaum, Rene Medema, and Helder Maiato for providing the cell line used, P. Todd Stukenberg for antibodies, and John R. Daum for technical assistance. This work was supported by grant 5R35GM126980 from the National Institute of General Medical Sciences.

## Supplemental Figures

**Figure S1.**
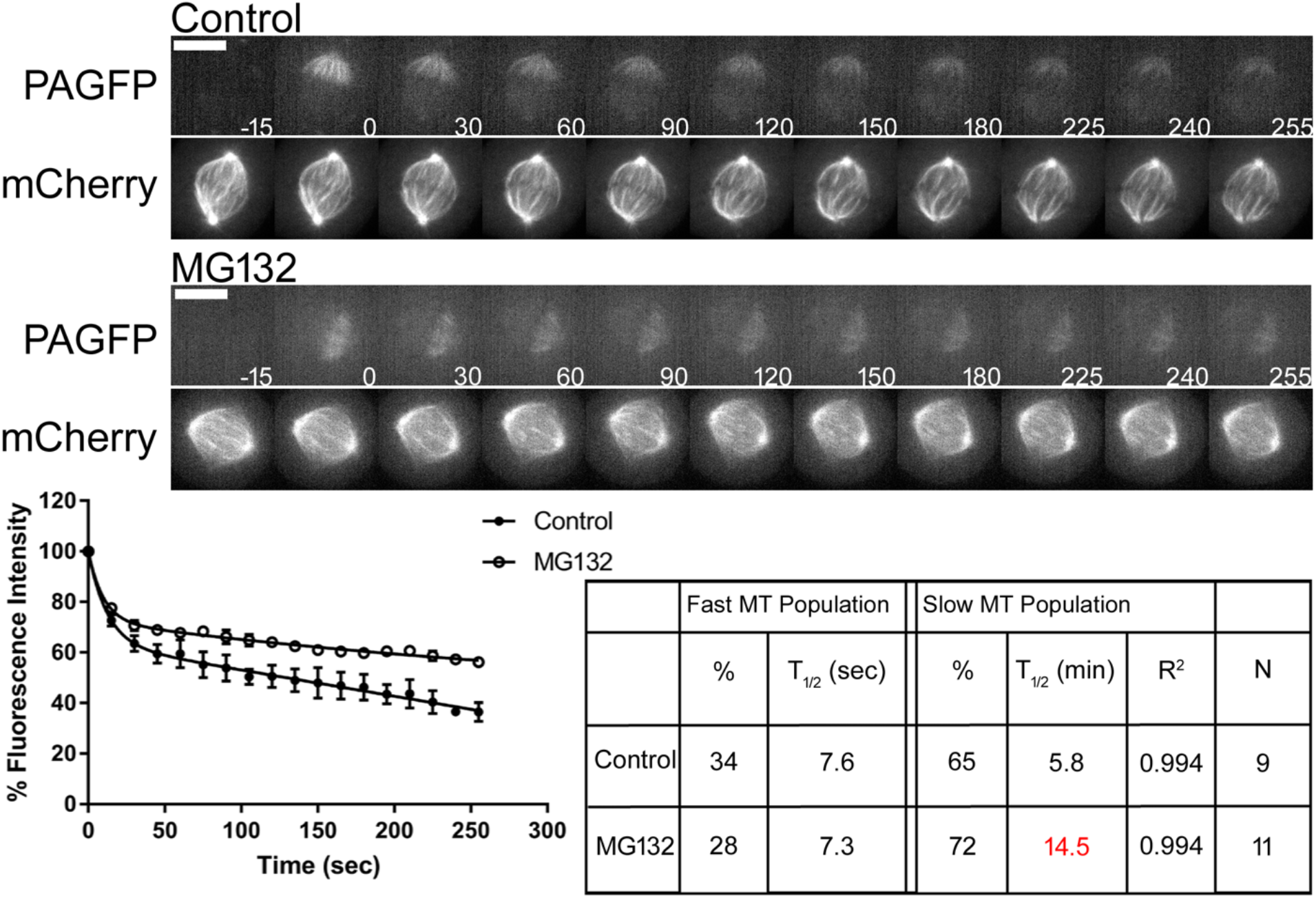
Prolonged metaphase arrest significantly decreases rate of turnover of slow microtubule population. (Top) Select frames from live cell imaging of Tubulin photoactivation in metaphase U2OS cells stably expressing photoactivatable GFP-Tubulin and mCherry-Tubulin following treatment with either control DMSO or 10 μm MG132. Cells treated with MG132 were incubated 3 h prior to imaging. mCherry-Tubulin frames are from the mid-volume plane. Control cells were imaged at metaphase. Time, sec. Bar, 10 μm. (Bottom) Fluorescence dissipation after photoactivation. The filled and unfilled circles represent the average values recorded at each time point after photoactivation. The bars represent SEM. Control n = 9 cells, MG132 n = 11 cells from two independent experiments. Lines indicate fitted curves (Control R^2^ = 0.994; MG132 R^2^ = 0.994). Significant differences from control indicated in red in table.

**Figure S2.**
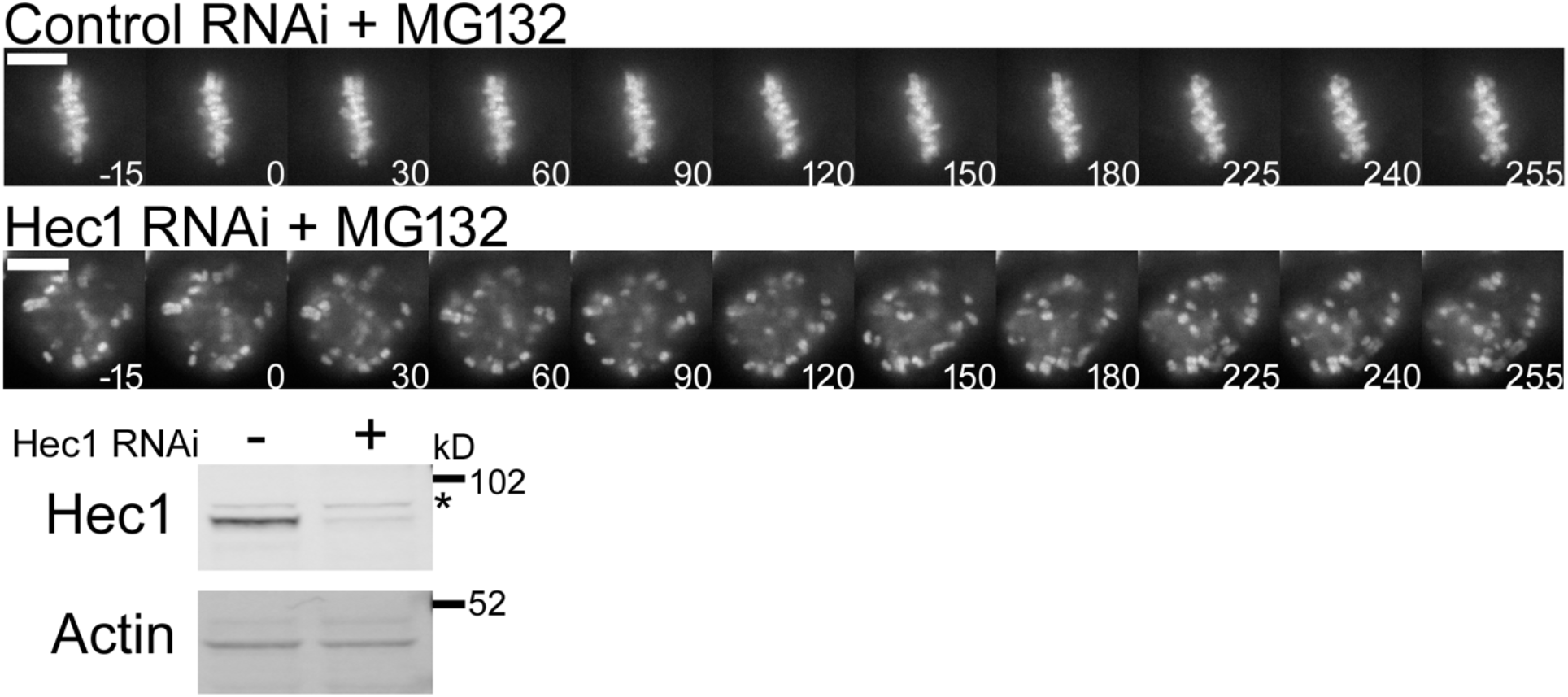
Hec1 depletion phenotype. (Top) Select mid-volume frames of DNA labeled with SiR-DNA from cells treated as in Fig. 1 demonstrating Hec1 depletion phenotype. Time, sec. Bar, 10 μm. (Bottom) Lysates from mitotic HeLa cells transfected with control or Hec1 siRNA and analyzed by Western blot probing for the indicated proteins.

**Figure S3.**
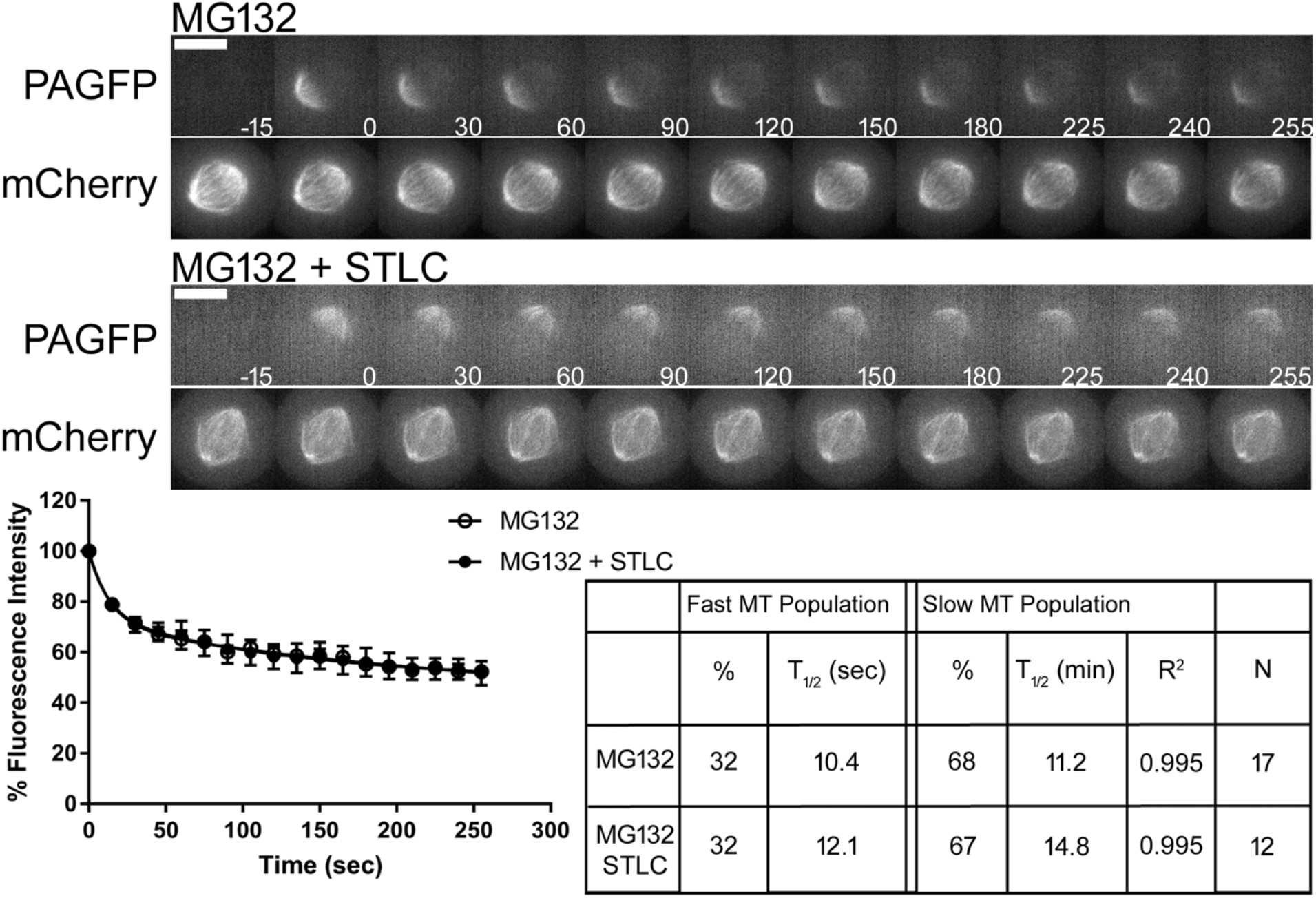
Eg5 inhibition does not collapse preformed bipolar spindles. (Top) Select frames from live cell imaging of Tubulin photoactivation in metaphase U2OS cells stably expressing photoactivatable GFP-Tubulin and mCherry-Tubulin following treatment with 10 μm MG132 for 3 h. Cells were then treated with either control DMSO or 10 μm STLC. Cells were incubated 30 min prior to imaging. mCherry-Tubulin frames are from the mid-volume plane. Time, sec. Bar, 10 μm. (Bottom) Fluorescence dissipation after photoactivation. The filled and unfilled circles represent the average values recorded at each time point after photoactivation. The bars represent SEM. MG132 n = 17 cells, MG132 + STLC n = 12 cells from four independent experiments. Lines indicate fitted curves (Control R^2^ = 0.995; MG132 + STLC R^2^ = 0.995).

**Figure S4.**
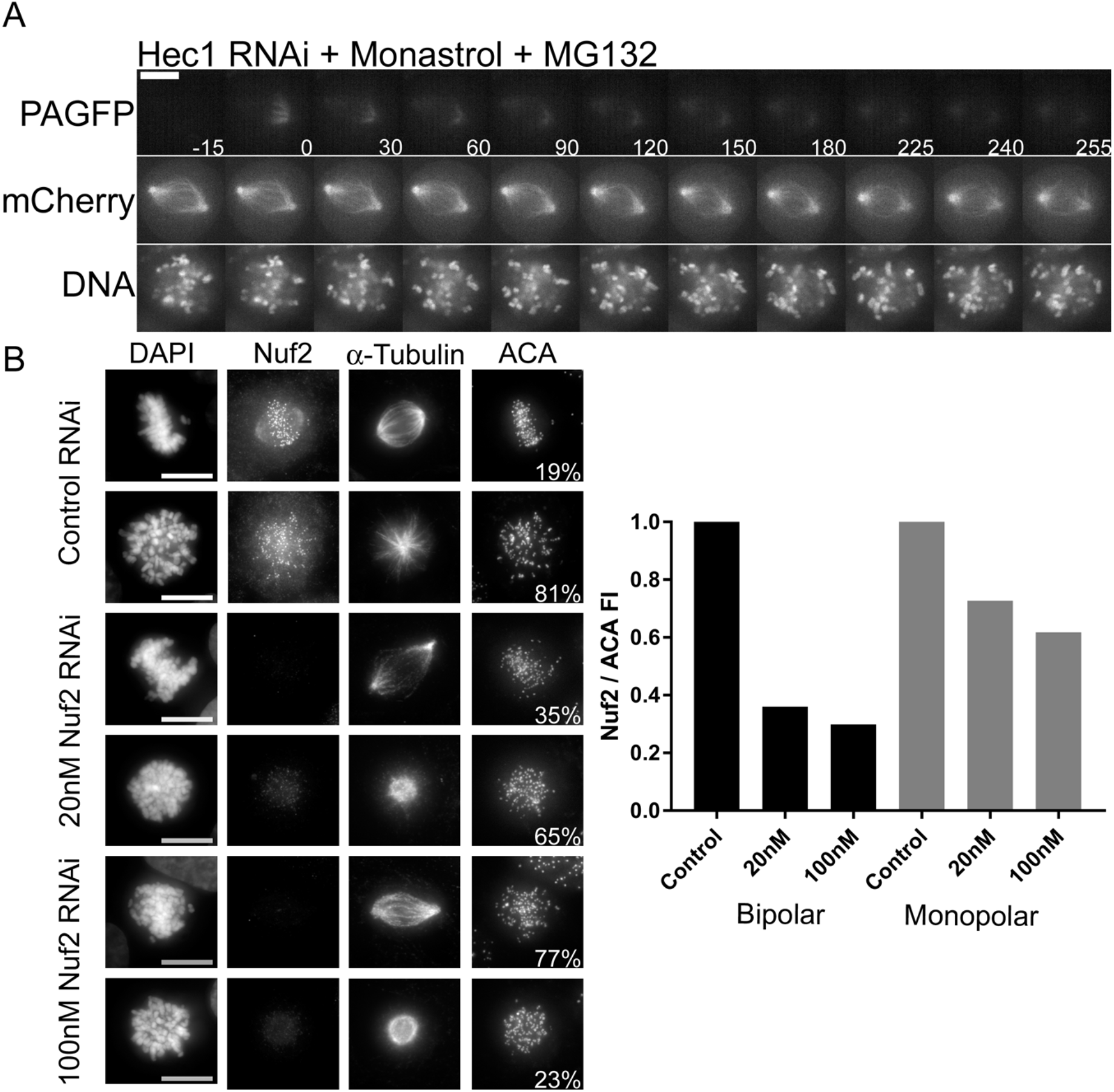
Maintenance of spindle bipolarity is sensitive to Hec1 and Nuf2 levels following Monastrol treatment. (A) Select frames from live cell imaging of Tubulin photoactivation in U2OS cells stably expressing photoactivatable GFP-Tubulin and mCherry-Tubulin following transfection with Hec1 siRNA along with 100 μm Monastrol and 10 μm MG132 treatment to prevent mitotic exit. mCherry-Tubulin and DNA labeled with SiR-DNA frames are from the mid-volume plane. Time, sec. Bar, 10 μm. (B, left) Immunofluorescence images of U2OS cells stably expressing photoactivatable GFP-Tubulin and mCherry-Tubulin immunostained for DNA (DAPI), Nuf2, α-Tubulin, and ACA following transfection of either control, 20 nM Nuf2, or 100 nM Nuf2 siRNA. Cells were treated with 100μm Monastrol for 1 h prior to fixation. n = 150 cells for each condition. Images represent maximum-intensity projections. Nuf2 images are scaled equivalently. Bar, 10 μm. (B, right) Fluorescence intensity quantification of Nuf2 levels from the indicated conditions in the left panel. n = 5-8 cells for each condition. FI, Fluorescence Intensity.

**Figure S5.**
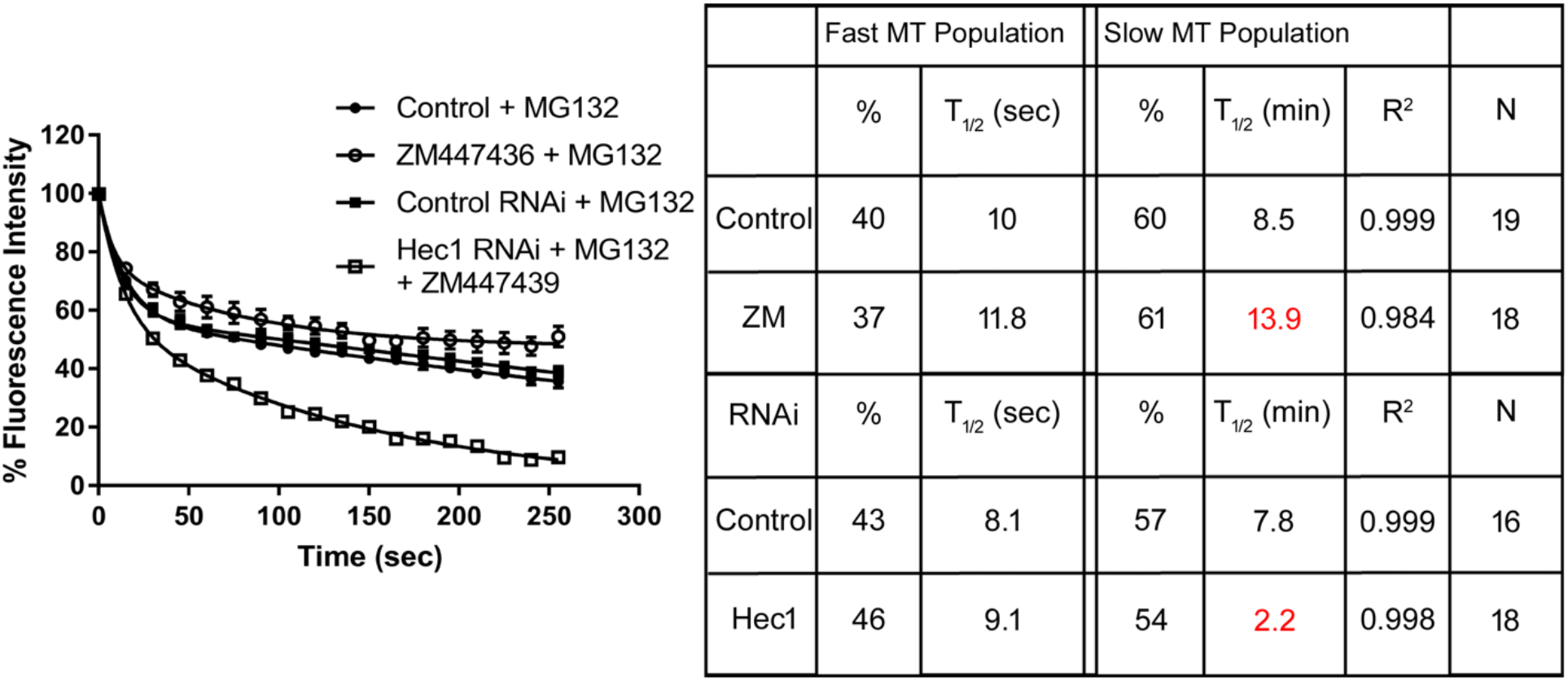
Compilation of data from Figure 4 and Figure 5. Aurora kinase inhibition decreases turnover of the slow population in cells with intact kinetochores but accelerates slow population turnover in cells where kinetochore-microtubule attachments are blocked.

**Figure S6.**
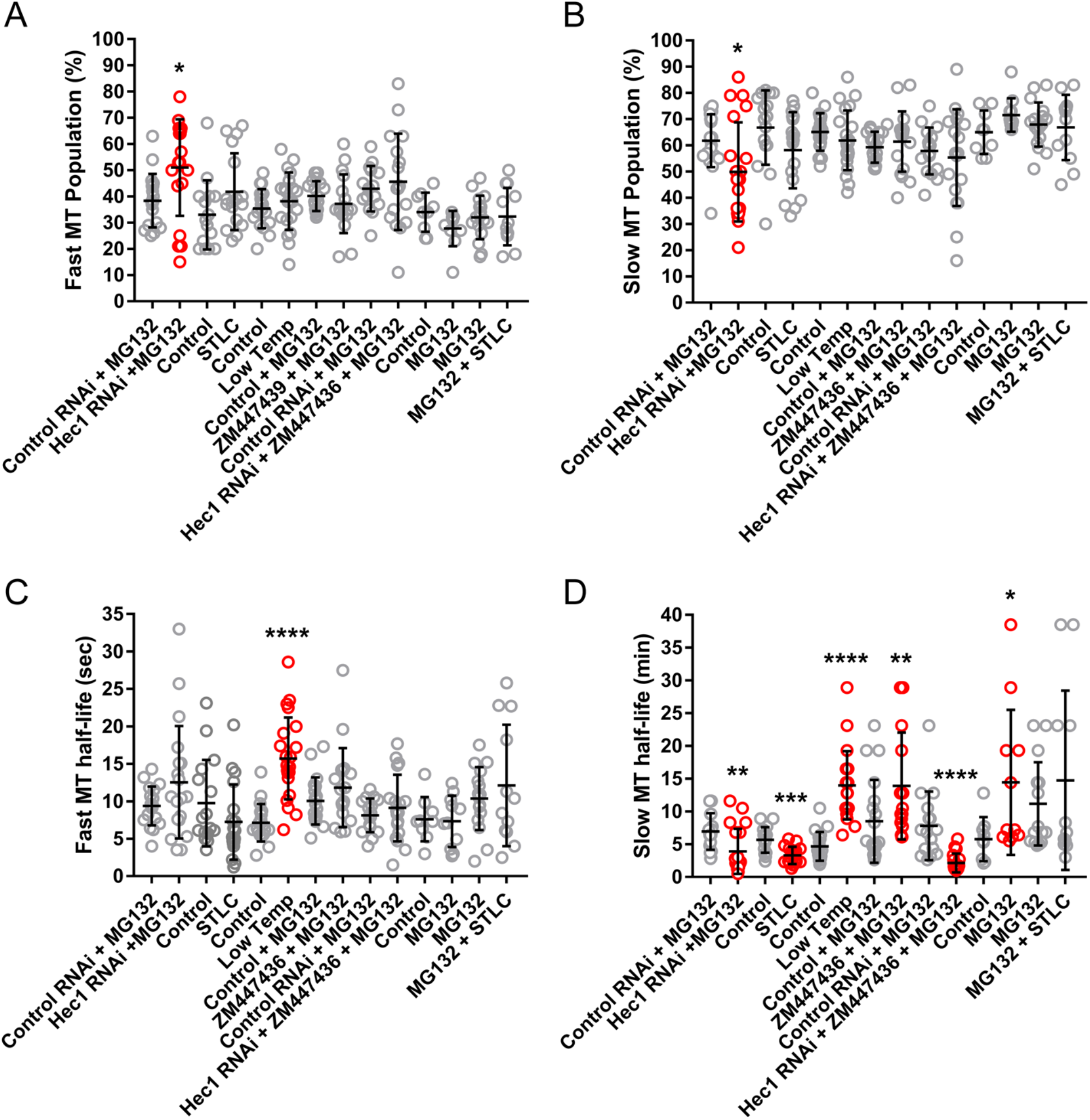
Compilation of total fluorescence dissipation after photoactivation data. (A) Fast microtubule population data. (B) Slow microtubule population data. (C) Fast microtubule turnover data. (D) Slow microtubule turnover data. *, P < 0.05; **, P < 0.01; ***, P < 0.001; ****, P < 0.0001. Statistical significance was determined by comparison to the appropriate experimental control and is indicated by red entries. Dots indicate data from individual cells. Lines show mean and SD.

## Video Legends

Movie S1. **Monopolar spindle formation in U2OS cell treated with 20 nM Nuf2 siRNA, monastrol and MG132.** Example of a cell that forms a monopolar spindle. The cell expresses photoactivatable GFP-Tubulin and mCherry-Tubulin and is labeled with Sir-DNA. For each time point, DNA and mCherry-Tubulin images were acquired. Total time is 42 min. Bar, 10 μm.

Movie S2. **Bipolar spindle formation in U2OS cell treated with 100 nM Nuf2 siRNA, monastrol and MG132.** Example of a cell that forms a bipolar spindle. The cell expresses photoactivatable GFP-Tubulin and mCherry-Tubulin and is labeled with Sir-DNA. For each time point, DNA and mCherry-Tubulin images were acquired. Total time is 42 min. Bar, 10 μm.

